# Steryl ester formation and accumulation in steroid-degrading bacteria

**DOI:** 10.1101/775916

**Authors:** Johannes Holert, Kirstin Brown, Ameena Hashimi, Lindsay D. Eltis, William W. Mohn

## Abstract

Steryl esters (SEs) are important storage compounds in many eukaryotes and are often prominent components of intracellular lipid droplets. Here we demonstrate that selected Actino- and Proteobacteria growing on sterols are also able to synthesize SEs and to sequester them in cytoplasmic lipid droplets. We found cholesteryl ester (CE) formation in members of the actinobacterial genera *Rhodococcus*, *Mycobacterium*, and *Amycolatopsis* as well as several members of the proteobacterial Cellvibrionales order. CEs maximally accumulated under nitrogen-limiting conditions, suggesting that steryl ester formation plays a crucial role for storing excess energy and carbon under adverse conditions. *Rhodococcus jostii* RHA1 was able to synthesize phytosteryl- and cholesteryl esters, the latter reaching up to 7% of its cellular dry weight and 69% of its lipid droplets. Purified lipid droplets from RHA1 contained CEs, free cholesterol and triacylglycerols. In addition, we found formation of CEs in *Mycobacterium tuberculosis* when grown with cholesterol plus an additional fatty acid substrate. This study provides a basis for the application of bacterial whole cell systems in the biotechnological production of SEs for use in functional foods and cosmetics.

**IMPORTANCE:** Oleaginous bacteria exhibit great potential for the production of high-value neutral lipids, such as triacylglycerols and wax esters. This study describes the formation of steryl esters (SEs) as neutral lipid storage compounds in sterol-degrading oleaginous bacteria, providing a basis for biotechnological production of SEs using bacterial systems with potential applications in the functional food, nutraceutical, and cosmetic industries. We found cholesteryl ester (CE) formation in several sterol-degrading Actino- and Proteobacteria under nitrogen limiting conditions, suggesting an important role of this process in storing energy and carbon under adverse conditions. In addition, *Mycobacterium tuberculosis* grown on cholesterol accumulated CEs in the presence of an additional fatty acid substrate.

## INTRODUCTION

Many bacteria synthesize and accumulate carbon and energy storage compounds under environmental stress conditions to increase their survival in environments with fluctuating nutrient conditions (1). A much smaller subset of bacteria are oleaginous and accumulate neutral lipids, namely triacylglycerols (TAGs) and wax esters (WEs), as storage compounds. Although TAG and WE synthesis is rarely described in other bacteria, their accumulation is widespread within the actinobacterial genera *Streptomyces*, *Nocardia*, *Rhodococcus*, *Mycobacterium*, *Dietzia*, and *Gordonia*, and has also been reported in the proteobacterial genera *Acinetobacter*, *Alcanivorax* and *Marinobacter* (2–7). TAGs and WEs are highly hydrophobic, non-polar lipids, which are formed by esterification of a glycerol molecule to three fatty acids or by esterification of a long chain fatty alcohol to a fatty acid, respectively (Fig. 1). Potential biotechnological applications of TAG and WE producing bacteria gained increasing attention over the last years due to the considerable industrial relevance of these compounds as biofuels and other products such as feed additives, cosmetics, and lubricants (8). Especially oleaginous *Rhodococcus* strains have been suggested as promising microbial cell factories for the large scale production of neutral lipids due to their ability to convert agroindustrial wastes into valuable lipids and their accessibility to cell engineering for increased lipid production (9).

**Fig. 1:**
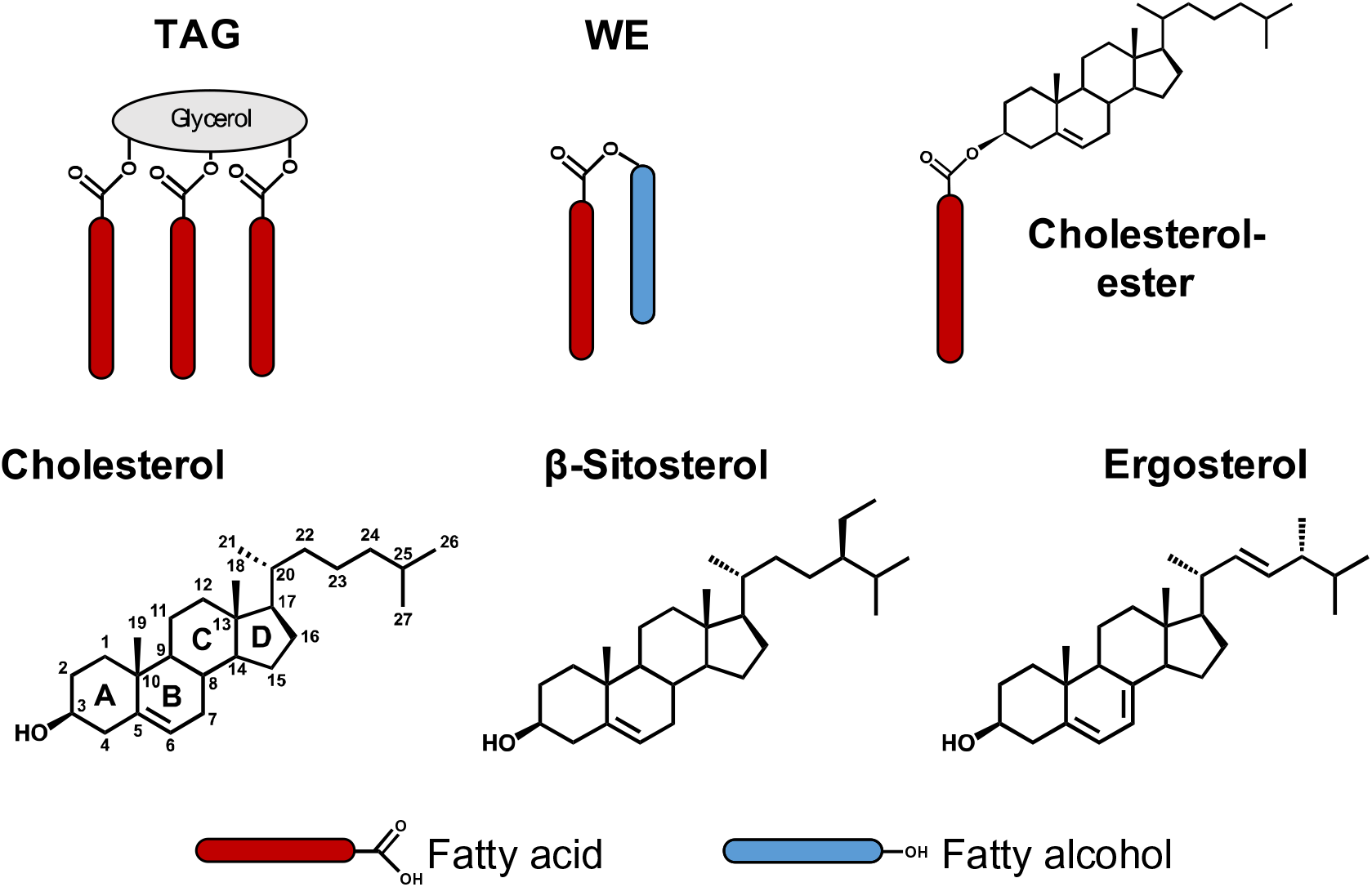
Schematic representation of triacylglycerols (TAGs) and wax esters (WEs) as neutral lipid storage compounds in bacteria and of a cholesterol-ester. Structures of the major animal, plant, and fungal sterols cholesterol, β-sitosterol, and ergosterol.

Accumulation of neutral lipids in bacteria is typically induced by environmental stress conditions, in which a suitable carbon source is in excess while additional growth is restricted by a deficiency of other nutrients or environmental conditions. The most prominent example is nitrogen limitation, which induces TAG and WE synthesis in a wide range of neutral lipid-accumulating Actino- and Proteobacteria. This has been most extensively studied in oleaginous *Rhodococcus* strains, which can accumulate up to 70% of their cellular dry weight as TAGs upon nitrogen depletion, depending on the available carbon source (10–12). In *Mycobacterium tuberculosis*, the causative agent of tuberculosis, dormancy inducing conditions such as nitric oxide (NO) stress or iron limitation have been shown to initiate accumulation of TAGs and WEs (13, 14). These intracellular carbon and energy storage compounds have been suggested to be important for the pathogenicity of *M. tuberculosis* (15, 16).

TAGs and WEs are sequestered into cytoplasmic lipid droplets, which are multifunctional organelles encapsulated by a phospholipid monolayer and specific proteins (17). Analogous to oleaginous bacteria, many eukaryotes also sequester neutral lipids as carbon and energy reservoirs into cytoplasmic lipid droplets. In contrast to bacterial lipid droplets, many eukaryotic lipid droplets contain sterols in addition to TAGs and WEs (18). Sterols constitute a key class of lipids in most animals, fungi, and plants, where they play essential roles as membrane constituents and precursors for vitamins and steroid hormones. The dominant sterol in animals is cholesterol, while plants have β-sitosterol and other phytosterols, and fungi have ergosterol (Fig. 1). Within eukaryotic lipid droplets, the sterol 3-hydroxyl group is typically esterified with fatty acids forming so-called steryl esters (SEs) (Fig. 1), which act as a sterol storage pool. Synthesis and breakdown of SEs plays a vital role in sterol homeostasis and imbalances between free and esterified cholesterol pools are related to several human diseases, such as Wolman disease, cholesteryl ester storage disease, atherosclerosis, and Alzheimer’s disease (19). In contrast to eukaryotes, sterol synthesis is rare in bacteria and mainly restricted to *Myxococcales* and *Methylococcales* (20). These bacterial sterols have structural features that distinguish them from eukaryotic sterols (21). Further, it is not known whether these bacteria are able to esterify and store their sterols in lipid droplets.

Sterols are abundant growth substrates for bacteria in the environment, and sterol degradation has been characterized in many actinobacterial isolates and is highly conserved in several genera of the Corynebacterineae suborder, namely the genera *Amycolicicoccus, Dietzia, Gordonia, Mycobacterium, Nocardia, Rhodococcus,* and *Tsukamurella* (22). In addition, we recently isolated several sterol-degrading strains belonging to the gammaproteobacterial order Cellvibrionales (23), which are also able to degrade sterols such as cholesterol. Under aerobic conditions, sterols are degraded by a conserved pathway into central metabolic moieties such as acetyl-CoA and propionyl-CoA (24), which are subsequently channeled into central metabolic pathways for energy conservation and production of biomass. Among those sterol-degrading bacteria are many that are also well known for their ability to produce TAGs and WEs as neutral lipid storage compounds, such as the Actinobacteria, *Rhodococcus jostii* RHA1 (RHA1) (25) and *Mycobacterium tuberculosis* (*Mtb*) (26). Key enzymes in bacterial TAG and WE synthesis are CoA-dependent, bifunctional WS/DGAT enzymes encoded by *atf* genes. These enzymes catalyze both the acetylation of DAGs to TAGs and of fatty alcohols to WEs (27). Most WS/DGATs are promiscuous enzymes accepting a variety of acyl-CoAs and alcohols as substrates (5, 28). Interestingly, the WS/DGAT enzyme AtfA from the oleaginous, non-sterol degrading Proteobacterium *Acinetobacter calcoaceticus* ADP1 has been shown to have SE-synthase activity when heterologously expressed in *E. coli* or the yeast *Saccharomyces cerevisiae* (29). Over 50 years ago, two studies reported that different *Mycobacterium* strains produced small amounts of cholesteryl fatty acid esters when incubated with cholesterol in cell suspensions (30) and that *Mycobacterium smegmatis* also produced esters of other sterols (31). However, these studies did not provide further insight into the cellular localization of SE accumulation or which incubation conditions elicit SE formation, and no further reports have been published on bacterial SE synthesis since then.

Here we investigate the ability of sterol-degrading bacteria to synthesize lipid storage compounds from sterol substrates under adverse growth conditions and report formation of SEs as lipid storage compounds in bacteria. This significantly extents the knowledge about bacterial SE formation and provides a basis for biotechnological production of SEs using bacterial systems with potential applications in the functional food, nutraceutical, and cosmetic industries.

## RESULTS

### Accumulation of SEs in RHA1 under nitrogen limiting conditions

To investigate formation of lipid-storage compounds from sterol substrates, *Rhodococcus jostii* RHA1 was grown in mineral medium with 1 mM cholesterol or 1 mM of a phytosterol mix, containing 76.4% β-sitosterol, 10.4% β-sitostanol, 7.3% campesterol and 1% campestanol, under nitrogen-limiting and nitrogen-excess conditions. In cholesterol-grown cultures, RHA1 reached a biomass in early stationary phase that was approximately four-fold higher under nitrogen excess than under nitrogen-limiting conditions (Fig. 2A). In contrast to the N-excess cultures, the N-limited cultures had no residual ammonium but did have residual cholesterol. Similar results were obtained during growth on the phytosterol substrate mix (not shown). Thin layer chromatography (TLC) analysis of neutral lipid extracts revealed compound spots with the same *R*_f_-value as an authentic cholesterol-palmitate standard (*R*_f_ = 0.78) in N-limited cholesterol and phytosterol cultures, but not in the respective N-excess cultures or in N-limited or N-excess cultures grown on non-sterol substrates (Fig. 2B). In addition, compounds with similar Rf-values as an authentic tripalmitin TAG standard (*R*_f_ = 0.24) were present in extracts of all N-limited cultures. Time-dependent sampling of RHA1 grown on cholesterol under N-excess and N-limiting conditions revealed that increased formation of the presumptive cholesteryl esters coincided with the depletion of ammonium from the medium (Fig. 3A). Under N-excess conditions only minimal amounts of these compounds accumulated during exponential growth and stationary phase (Fig. 3B). When nitrogen was replenished in an N-limited, stationary-phase, cholesterol-grown culture, the accumulated lipids including the presumptive SEs disappeared within 72 h (Fig. 2B).

**Fig. 2:**
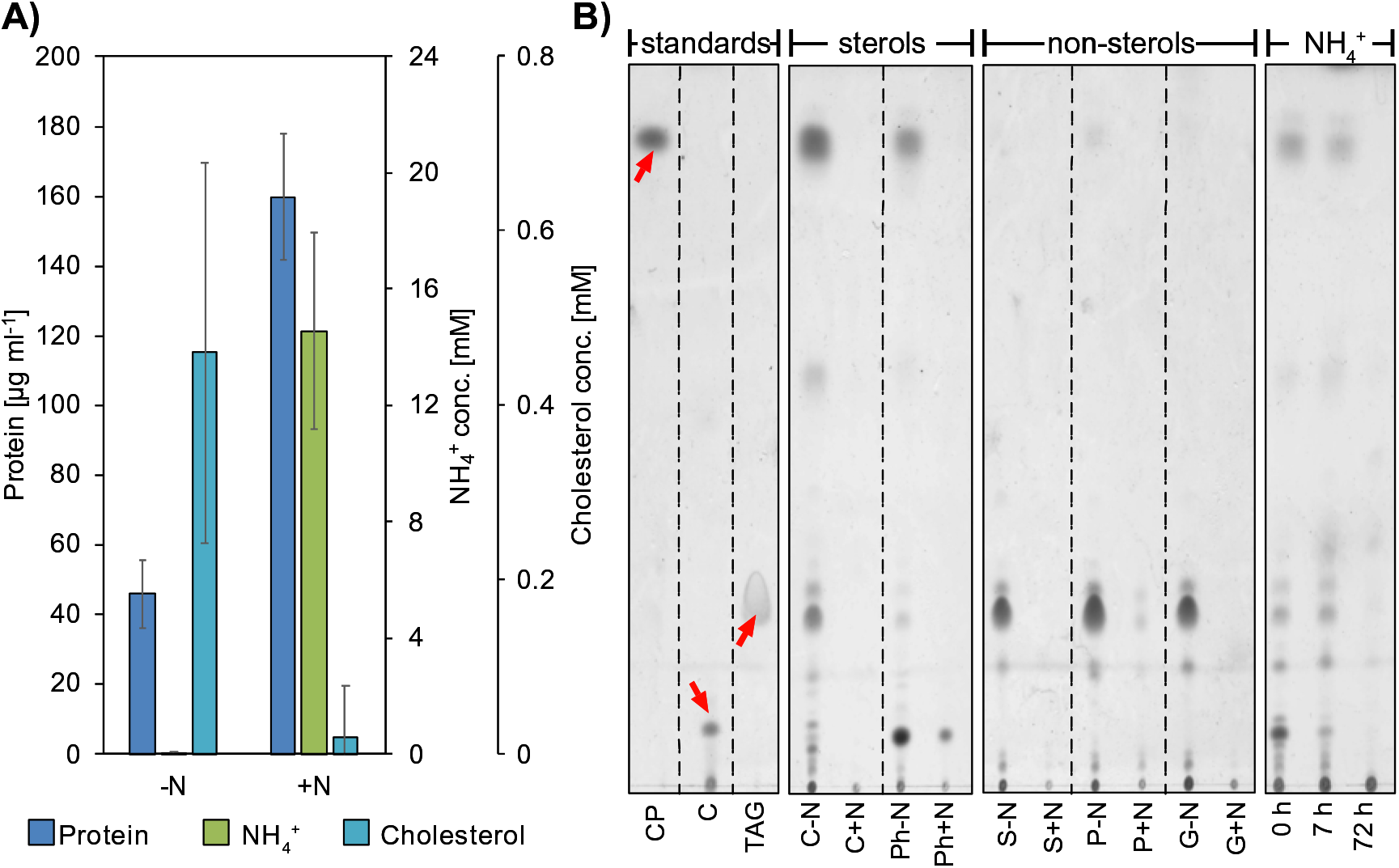
Growth of strain RHA1 under nitrogen-limiting and nitrogen-excess conditions. **A)** at early stationary phase (around 100 h). **B)** TLC analysis of neutral lipid extracts from early stationary phase N-limited (-N) and N-excess (+N) cultures grown with different carbon sources. Abbreviations: CP, cholesterol palmitate; C, cholesterol; TAG, triacylglycerols; Ph, phytosterol mix; S, succinate; P, pyruvate; G, glucose; NH_4_^+^, N replenished in stationary-phase C-N culture. The volume of neutral lipid extracts applied to TLC plates was normalized on the extracted cellular dry weight of each culture. Red arrows indicate standard spots. Results were obtained on two TLC plates which were treated exactly the same way throughout the experiment. The results of one representative experiment out of several biological and technical replicates are shown for each substrate.

**Fig. 3:**
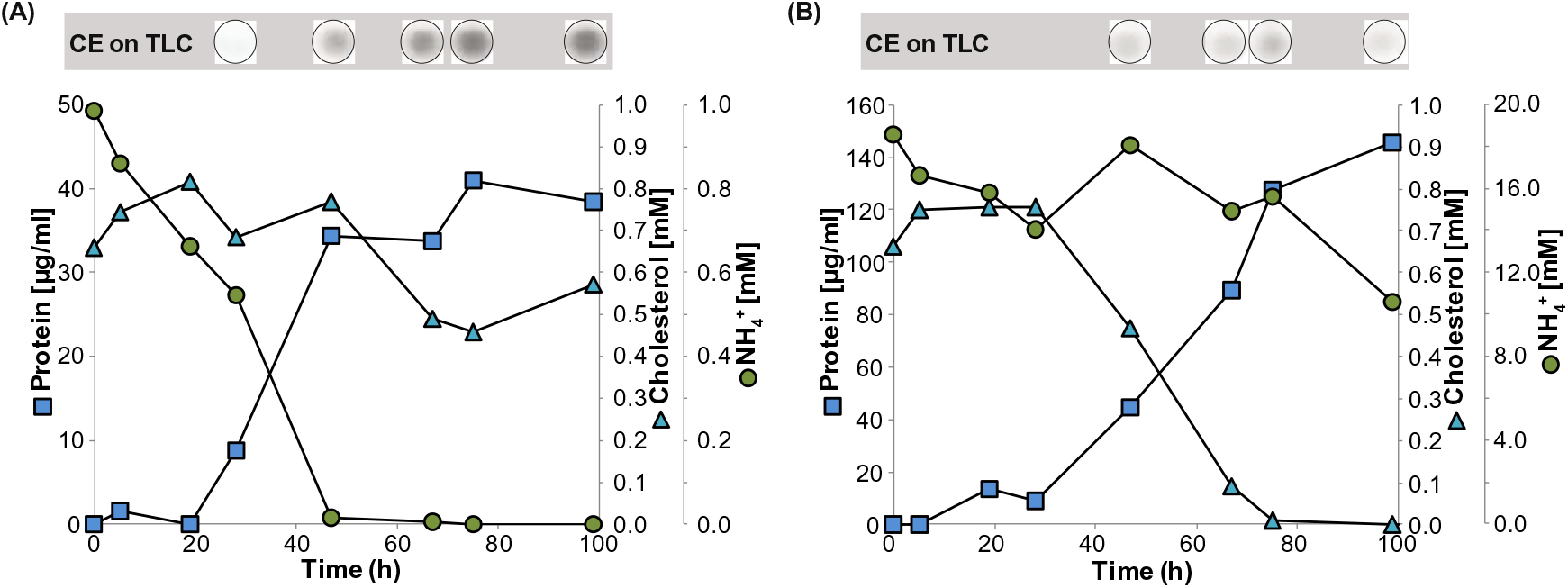
Growth of RHA1 on 1 mM cholesterol under nitrogen-limiting conditions **(A)** and nitrogen-excess conditions **(B)**. The volume of neutral lipid extracts applied to TLC plates was normalized on the basis of culture biomass as protein. One of two reproducible experiments is shown.

To confirm the formation of CEs, we analyzed native neutral lipid extracts from strain RHA1 grown with cholesterol or succinate under N-limiting conditions in early stationary phase using a gas chromatography (GC)-coupled flame ionization detector (FID). Comparison to a cholesteryl-palmitate (C16:0) and a cholesteryl-heptadecanoate standard (C17:0) confirmed that these CEs accumulated in cholesterol-grown RHA1 cells (Fig. 4A), but were absent in succinate-grown cells. In addition, five other compounds from cholesterol-grown cells eluted close to these CEs, which were absent in succinate-grown cells, suggesting that seven CE species, with different fatty acid chain lengths, are formed with cholesterol as substrate. Comparison to a tripalmitate ((C16:0)_3_) TAG standard suggested the accumulation of up to nine different TAG species in strain RHA1 grown with cholesterol and eleven different TAG species in cells grown with succinate (Fig. 4A). Time-dependent sampling starting at the onset of nitrogen depletion from the media followed by GC-FID analysis showed that the CE:TAG ratio shifted from approximately 60% to 72% within the first 24 h of incubation and to 69% after 48 h.

**Fig. 4:**
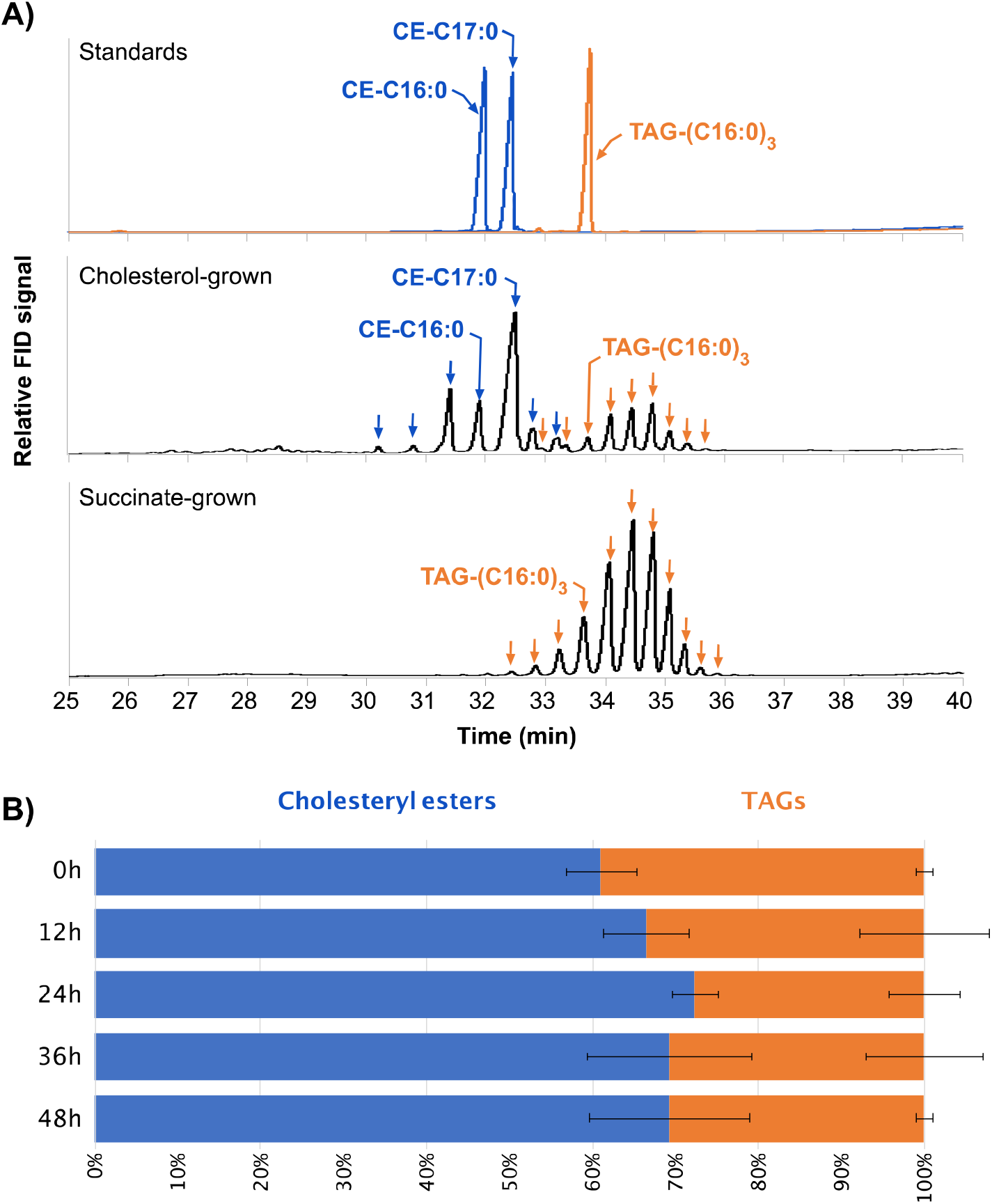
GC-FID analysis of accumulating neutral lipids in nitrogen-limited RHA1 cultures grown with cholesterol or succinate. **(A)** GC-FID chromatograms of authentic cholesteryl palmitate (C16:0) and cholesteryl heptadecanoate (C17:0) standards (blue line) and of a trimpalmitin TAG (C16:0_3_) standard (orange line) and of neutral lipid extracts of cholesterol- and succinate-grown cells harvested 24 h after nitrogen depletion. Seven accumulating CE species were identified either by comparison to the standards or based on the proximity of their respective peaks to the standard peaks and their absence in succinate grown cells. Identified and predicted CEs are labelled with blue arrows, identified and predicted TAG species are labelled with orange arrows. **(B)** Ratio of accumulating CEs (blue bars) and TAGs (orange bars) in cholesterol-grown nitrogen-limited RHA1 cultures over time starting at nitrogen depletion (0 h). Peak areas of the seven CE- and nine TAG-peaks shown in Fig. 4A were normalized on the peak area of the internal standard cholestane (not shown) before calculating their respective sums. Average values of three biological replicates and their standard deviations are shown.

To gain further insight into the nature of SEs accumulating in strain RHA1, neutral lipid extracts of cholesterol- and phytosterol-grown cultures were fractionated on silica columns producing five lipid fractions (**supplemental Fig. S1**). SEs eluted in fraction 2 and were well separated from other lipids, like TAGs (Fractions 3 and 4) and free sterols (fractions 5). Aliquots of fraction 2 were subjected to alkaline transesterification with methanol. GC-MS analysis showed that free cholesterol or the free phytosterols (β-sitosterol, β-sitostanol, and campesterol) were formed from transesterification, which were not present in untransesterified extracts (Fig. 5A-B). Only the respective sterols provided as a growth substrate were detected in this analysis. In addition, the lack of other free alcohols after transesterification confirmed the absence of wax esters in neutral lipid extracts. In addition, a range of seven fatty acid methyl esters (FAMEs) were detected after transesterification of cholesteryl esters, supporting the results from the GC-FID analysis. Those were identified as methyl esters of C15:0, C16:1, C16:0, C17:1, C17:0, C18:1 and C18:0 (Fig. 5A). The most abundant fatty acids were the odd-numbered fatty acids, C17:0, C17:1, and C15:0, which constituted more than 75% of produced FAMEs (Fig. 5C). GC-MS analysis of transesterified fraction 2 confirmed that no detectable amounts of other free alcohols indicative of wax esters were present (not shown). Analysis of all neutral lipid extracts using a GC-MS method for wax-ester analysis (32) showed that no detectable amounts of wax esters were present in any sample (not shown), which is in agreement with earlier reports that wax esters are only formed in minimal amounts in RHA1 (32).

**Fig. 5:**
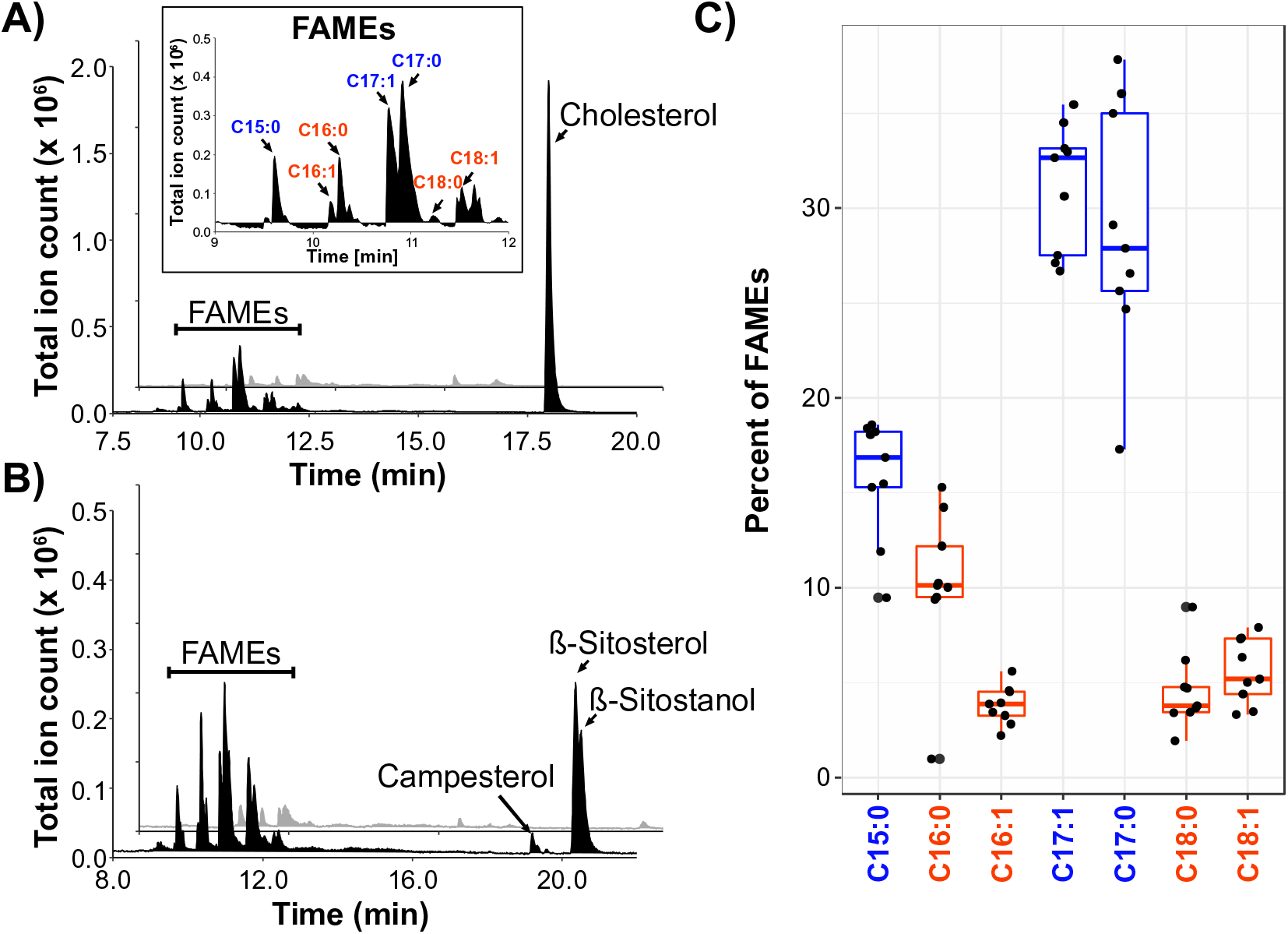
Identification of SEs accumulating in N-limited RHA1 cultures. GC-MS analysis of transesterified fraction 2 (black peaks) and untreated fraction 2 (grey peaks) of neutral lipid extracts of nitrogen-limited cultures grown with **(A)** cholesterol and **(B)** phytosterols. Transesterification of SEs produced free cholesterol and the free phytosterols β-sitosterol, β- sitostanol, and campesterol, as well as seven fatty acid methyl esters of C15:0, C16:1, C16:0, C17:1, C17:0, C18:0 and C18:1. One representative GC-MS chromatogram is shown. **(C)** Fatty acid profile of transesterified fraction 2 derived from nine cholesterol-grown cultures grown under nitrogen-limiting conditions from five independent experiments harvested approximately 24h after nitrogen depletion. Odd-numbered fatty acids are coloured blue and even-numbered fatty acids are coloured red

This confirmed that the previously unidentified lipids accumulating in N-limited RHA1 cultures grown with cholesterol or phytosterols are fatty acid steryl esters. In the late stationary phase (100 h) CEs accounted for 6.8 +/− 1.4% of the cellular dry weight in N-limited cultures (n = 5 from two independent experiments) based on GC-MS quantification.

### Localization of accumulated cholesterol and CEs in RHA1

To confirm that accumulating neutral lipids are localized intracellularly, we stained stationary phase, cholesterol-grown, washed, N-excess, and N-limited RHA1 cells with the lipophilic fluorescence stain Nile Red. Microscopic analysis showed that stained lipids accumulated as lipid droplets in the cell interior (**supplemental Fig. S2**) of N-limited cells but were absent from N-excess cells. In addition, we incubated RHA1 with fluorescently labelled cholesterol (BODIPY-cholesterol) and sodium palmitate for 48 h without a nitrogen source. Microscopic analysis showed that fluorescently labelled cholesterol was mainly present within lipid droplets inside the cells (**supplemental Fig. S3**) but mostly absent from cell membranes. TLC analysis of neutral lipid extracts of cells with fluorescently labelled cholesterol showed that no CEs were formed (not shown), which is most likely because the altered chemical structure of the labelled cholesterol side chain inhibits esterification. These results confirm that cholesterol and CEs are mainly stored in intracellular lipid droplets.

To gain further insight into the location of CEs within RHA1 cells, lipid droplets were isolated from RHA1 cultures grown with cholesterol or sodium palmitate or a mixture of both substrates under N-limiting conditions. With all three substrate combinations, gradient centrifugation of total cell extracts produced visible bands of lipid droplet on top of the sucrose gradient (**supplemental Fig. S4**). TLC analysis revealed that CEs and free cholesterol were present in neutral lipid extracts of lipid droplets derived from cells grown on cholesterol but not in lipid droplets derived from cells grown on palmitate alone, while TAGs were present under all three conditions (Fig. 6A). This was confirmed by fractionation and subsequent transesterification of neutral lipid extracts (not shown). In addition, the lack of other free alcohols after transesterification confirmed the absence of wax esters in lipid droplets. In two independent experiments 52% and 58% of lipids in lipid droplets derived from cells grown on cholesterol were identified as CEs, 31% and 15% as TAGs and 17% and 27% as free cholesterol. Similar results were obtained for lipid droplets derived from cells grown on cholesterol plus palmitate, with 50% and 69% CEs, 35% and 23% TAGs and 15% and 8% cholesterol, while TAGs were the only neutral lipids in lipid droplets derived from cells grown on palmitate (Fig. 6B). In lipid droplets from cells grown on cholesterol, between 60% to 70% of fatty acids in CEs and TAGs were odd-numbered (C15 and C17), with C17:0 and C17:1 making up around 90% of these fatty acids (Fig. 6C). In lipid droplets from cells grown with cholesterol plus palmitate, 23% to 55% of fatty acids were odd-numbered. TAGs of cells grown on palmitate contained almost exclusively (∼95%) even-numbered fatty acids. (Fig. 6C).

**Fig. 6:**
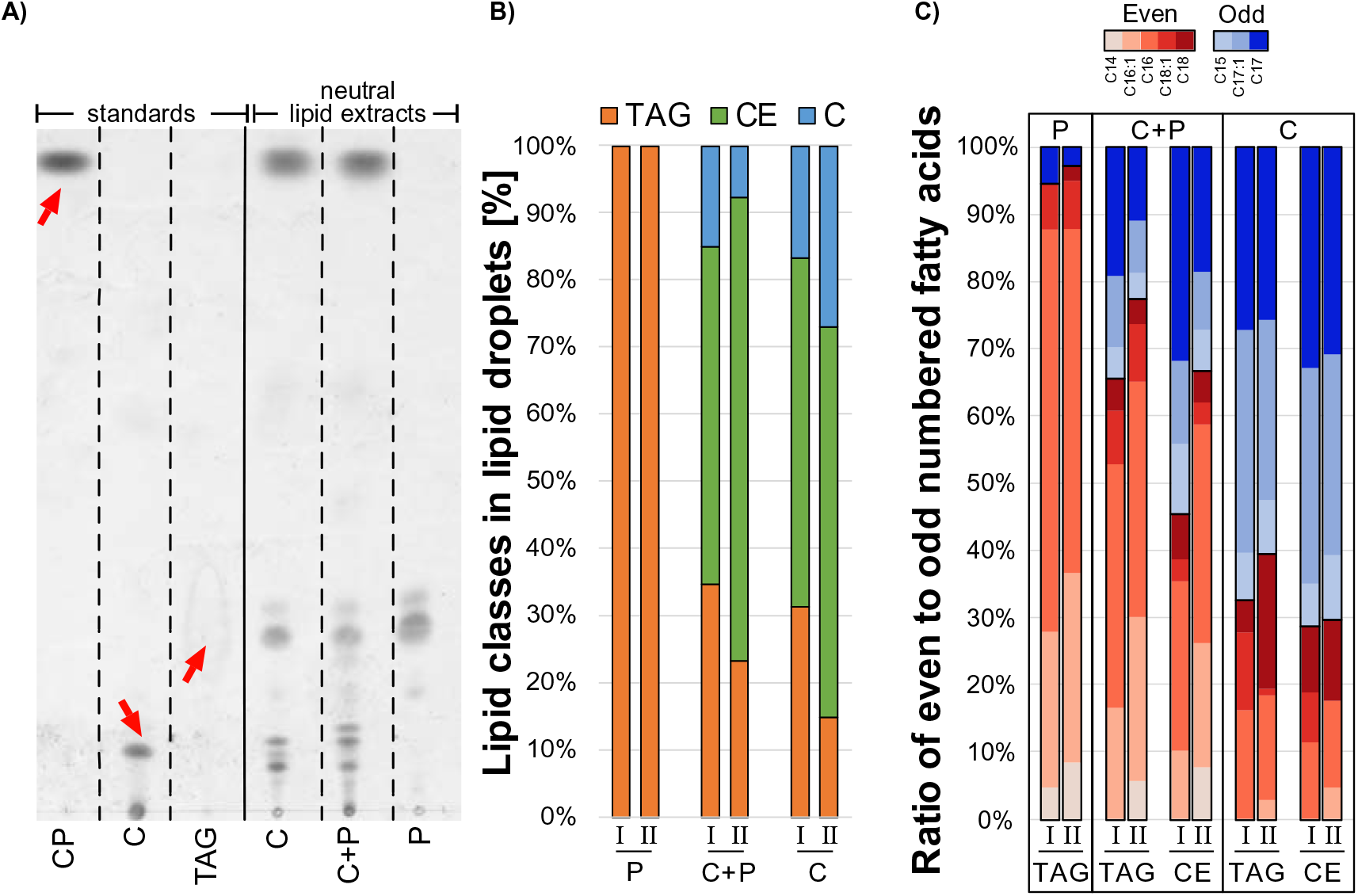
Isolation of lipid droplets from strain RHA1 grown with cholesterol (**C**) or palmitate (**P**) or a mixture of cholesterol plus palmitate (**C+P**) under nitrogen-limiting conditions. **A)** TLC analysis of neutral lipids derived from total lipid extracts from isolated lipid droplets. Abbreviations as in previous figures. **B)** Molar ratio of TAGs, CEs and free cholesterol in lipid droplets derived from cholesterol, cholesterol plus palmitate and palmitate grown cells. **C)** Ratio of even to odd-numbered fatty acids in CEs and TAGs in lipid droplets and the respective contributing fatty acid chain lengths. The results of two independent experiments (I and II) are shown in **B)** and **C)** and of experiment II in **A)**.

### Formation of SEs in other sterol-degrading bacteria

To test whether steryl ester formation also occurs in other sterol-degrading bacteria, we extracted neutral lipids from the Actinobacteria, *Mycobacterium smegmatis* strain mc^2^155, *Mycobacterium* sp. strain BC8-1, *Rhodococcus opacus* strain Chol-4, *Rhodococcus ruber* strain Chol-4, and *Amycolatopsis* sp. strain ATCC 39116, as well as the Gammaproteobacteria, *Zhongshania* sp. strain SB11-1A, *Haliaceae* sp. strain BC5-1, and the BD1-7 clade bacterium strain SB11-3 grown on cholesterol under nitrogen-limiting and nitrogen-excess conditions. The latter strains were isolated in our lab (23) and are among the first Proteobacteria reported to be able to degrade sterols. TLC analysis indicated these strains produced SEs in N-limited cultures in the stationary phase in differing amounts, with highest yields in the *Rhodococcus*, *Mycobacterium*, and *Zhongshania* strains and lower yields in *Amycolatopsis*, *Haliaceae*, and the BD1-7 clade strain (**supplemental Fig. S5A**). Subsequent fractionation of neutral lipid extracts followed by transesterification and GS-MS analysis confirmed the formation of CEs in all strains (**supplemental Fig. S5B**).

In addition, we tested steryl ester formation in *Mycobacterium tuberculosis* (*Mtb*) Erdmann growing on cholesterol and on cholesterol plus sodium palmitate. Because *Mtb* did not grow when sodium palmitate was added to the medium prior to inoculation we spiked palmitate into cultures growing on cholesterol after 5 days. TLC analysis of neutral lipid extracts of *Mtb* indicated the formation of small amounts of CEs in cells growing on cholesterol after 7 days and in cells growing on cholesterol plus palmitate after 12 and 13 days, when cells were in stationary phase (**supplemental Fig. S6A-B**). Subsequent fractionation of neutral lipid extracts followed by transesterification and GS-MS analysis confirmed the formation of CEs (**supplemental Fig. S6C**). The only detectable FAME in this fraction was C16:0. Formation of free hexacosanol after transesterification suggested that a C_26_-OH wax ester was also formed. In addition, we tested whether nitric oxide (NO) stress or iron limitation, which are known inducers of neutral lipid accumulation in *Mtb* (13, 14), increase CE formation from cholesterol or cholesterol plus palmitate. For this, cholesterol-grown cell suspensions of *Mtb* were incubated with cholesterol alone or cholesterol plus palmitate. To induce NO stress, two pulses of 0.5 mM spermine-NONOate were added at the beginning of the experiment and again after 16 h of incubation and samples were taken after 16 h and after 2 days of incubation. Spermine (0.5 mM) was added to control cultures. Deferoxamine (0.5 mM) was added as an iron chelator and samples were taken after 2 and 4 days of incubation. Confirming the results from the initial growth experiment, all extracts from *Mtb* cells incubated with cholesterol plus palmitate contained CEs with C16:0 as only detectable FAME. However, addition of NO or deferoxamine did not increase the amount of accumulating CEs (**supplemental Fig. S6D**). No significant accumulation of CEs occurred in treated cells incubated with only cholesterol. Small amounts of free cholesterol were detected after transesterification of extracts of cell suspensions treated with NO and incubated for 2 days with cholesterol (not shown).

## DISCUSSION

Steryl esters (SEs) are important storage compounds in many eukaryotes and are often prominent components of intracellular lipid droplets. Here we demonstrate that selected sterol-degrading Actino- and Proteobacteria are also able to synthesize SEs and to sequester them in cytoplasmic lipid droplets. We found SE formation in all sterol-degrading strains investigated in this study, including members of the actinobacterial genera, *Rhodococcus*, *Mycobacterium*, and *Amycolatopsis,* as well as members of the proteobacterial Cellvibrionales order. This is supported by findings from the 1960s, which showed that some *Mycobacterium* strains were able to produce different SE in small amounts (up to 0.5 % of the cellular dry weight for cholesterol) (30, 31). The ability of other oleaginous bacteria to synthesize SEs has been overlooked in other studies, which exclusively used non-steroidal substrates, such as glucose, gluconate, or benzoate, when investigating neutral lipid accumulation. Our results suggest that the ability to accumulate excess sterols as an energy and carbon source under adverse conditions plays a crucial role in sterol-degrading bacteria.

Using *Rhodococcus jostii* RHA1 as an oleaginous, sterol-degrading model organism, we found accumulation of minor amounts of CEs during exponential growth on cholesterol when ammonium was available, and a significant increase in CE and TAG accumulation, when ammonium was limiting, indicating that nitrogen limitation stimulates SE formation. This extends previous reports that nitrogen limitation is a major inducer for the accumulation of TAGs and WEs in RHA1 with non-steroidal substrates (12, 33). While we found significant accumulation of CEs in the two free-living *Mycobacterium* strains mc^2^115 and BC8-1 under nitrogen limiting conditions with cholesterol as the only substrate, the pathogenic strain *Mycobacterium tuberculosis* (*Mtb*) Erdmann did not grow on cholesterol in nitrogen-limited medium. However, small amounts of cholesterol-palmitate accumulated in this strain when growing on cholesterol plus palmitate as substrates without any stress conditions, demonstrating that pathogenic *Mtb* strains are able to accumulate CEs when a sterol and a fatty acid are available. During infection, *Mtb* triggers formation of foamy macrophages (34, 35), which provide a cholesterol and fatty acid-rich microenvironment, and *Mtb* metabolizes these host-derived lipids during infection and persistence (36–38). Interestingly, cholesterol and fatty acid uptake are co-regulated in *Mtb* (39), suggesting that *Mtb* has simultaneous access to both lipid classes *in vivo*. SE synthesis and accumulation could act as a buffer system for excess lipids in *Mtb* providing a way to regulate intracellular lipid metabolism. In addition, SEs could serve as a carbon and energy reservoir in *Mtb* under adverse conditions. In a similar manner, TAG accumulation and sequestration in cytoplasmic lipid droplets was shown to occur *in vivo* in pathogenic *Mtb* (16, 40) and was proposed to be important for persistence and reactivation (41). Accumulation of TAGs and WEs was shown to be stimulated by various dormancy-inducing stress conditions, such as hypoxia, NO stress, iron limitation, and increased acidity, which induce expression of several WS/DGAT encoding genes in *Mtb* (13, 42–44). Although we did not detect increased formation of CEs in *Mtb* under NO stress or iron limiting conditions, a combination of stress factors as described in (42) might provide deeper insight into the regulation of SE synthesis in *Mtb*.

Lipid droplets have been identified in most domains of life, including many eukaryotes, as well as archaeal and bacterial prokaryotes (45). While the general structure of lipid droplets is conserved in all these organisms, comprising a neutral lipid core coated by a phospholipid monolayer and various peripheral proteins, the neutral lipid composition varies significantly between organisms (17, 46). Our results show that CEs, in addition to TAGs and free cholesterol, are sequestered in intracellular lipid droplets in RHA1. The lipid droplet composition in RHA1 grown on cholesterol is very similar to that of lipid droplets found in selected mammalian lipid droplet-producing cell lines (chinese hamster ovaries CHO K2, rat kidney normal NKR and immortalized human fibroblasts SV589) (47), with around 50% CEs, 30% TAGs and 20% free cholesterol. The sequestration of cholesterol in lipid droplets is in agreement with an earlier report showing that fluorescently labeled cholesterol predominantly localizes in lipid droplets in mycobacteria (48).

Interestingly, the majority of fatty acids stored in TAGs and CEs in cholesterol-grown RHA1 cells were odd-numbered fatty acids, with C17:0 and C17:1 as main fatty acids, which is in stark contrast to the mostly even-numbered fatty acids typically found in eukaryotic neutral lipids. Complete metabolism of one cholesterol molecule through the canonical bacterial 9,10-seco steroid-degradation pathway is proposed to yield four propionyl-CoAs, four acetyl-CoAs, one pyruvate and one succinyl-CoA (24). The preponderance of odd-chain fatty acids in neutral lipids accumulating in RHA1 with cholesterol as carbon source suggests that propionyl-CoA was a significant building block for neutral-lipid fatty acids. Elevated ratios of odd-numbered to even-numbered fatty acids have also been found in TAGs accumulating in RHA1 growing with benzoate under nitrogen limiting conditions, where propionyl-CoA was presumably generated through the methylmalonyl-CoA pathway from succinate (12). Cholesterol-derived propionyl-CoA has been shown to be toxic to *Mtb* if not metabolized (49), and incorporation of propionyl-CoA into cell wall lipids (50) and TAGs (51) has been suggested as a mechanism to detoxify propionate in *Mtb*. Cholesterol-degrading bacteria, such as RHA1 and *Mtb*, might encounter increased cellular propionyl-CoA levels under adverse growth conditions and the formation of odd-numbered fatty acid CEs might provide an additional sink for this toxic end-product. However, we did not detect odd-numbered fatty acid CEs in *Mtb*, which accumulated CEs in significant amounts only in the presence of an additional fatty acid substrate under the tested conditions.

Sterols are abundant growth substrates for bacteria in the environment and sterol degradation is conserved in many actinobacterial genera, particularly in members of the *Corynebacterineae* (22). Recently, we have also identified sterol-degradation in some isolates of the oligotrophic marine Gammaproteobacteria (OMG) group belonging to the Cellvibrionales order (23). Steroid-degrading bacteria are globally distributed and actinobacterial ones are particularly abundant within soil and rhizosphere as well as in deep ocean habitats. Our results indicate that SE formation is conserved in sterol-degrading bacteria, suggesting that this trait might play important roles in these environments. Supporting this hypothesis, most neutral-lipid accumulating Actinobacteria isolated from soil samples (52) belong to taxa in which the ability to degrade sterols is conserved (22) and are thus likely to produce SEs. *De novo* synthesis of sterols is rare in bacteria, being largely restricted to *Myxococcales* and *Methylococcales* (20) and it is not known whether these bacteria are able to store these sterols as esters.

We searched the genomes of strain RHA1 and of *Mtb* Erdmann for homologs of one prokaryotic and fifty eukaryotic proteins with characterized steryl ester synthase activity available in the Swiss-Prot database, including phosphatidylcholine-sterol *O*-acyltransferases (EC 2.3.1.43), diacylglycerol *O*-acyltransferases (EC 2.3.1.20), sterol *O*-acyltransferase (EC 2.3.1.26), and phospholipid:diacylglycerol acyltransferases (EC 2.3.1.158). None of these proteins had any significant similarity (accepted coverage > 85%, percent identity > 30%, E-values < 0.01) to RHA1 or *Mtb* proteins. Based on the multifunctionality and substrate promiscuity of TAG- and WE-producing WS/DGAT enzymes, it is feasible that these enzymes might carry out an additional function as steryl ester synthases. Out of the 16 predicted WS/DGATs in RHA1, Atf6 (WP_011594556.1) and Atf8 (WP_011597548.1) have the highest amino acid sequence identity (40% and 39%, respectively) to AtfA from the oleaginous, non-sterol degrading Proteobacterium *Acinetobacter calcoaceticus* ADP1, which was shown to have SE-synthase activity when heterologously expressed in *E. coli* or *S. cerevisiae* (29). TAG and WE synthase activities have been confirmed for Atf8 and Atf6, respectively (12, 28) as well as for an Atf6 ortholog in *Rhodococcus opacus* PD630 (53), and deletions of the respective genes significantly decreased neutral lipid accumulation. In addition, transcription of *atf8* was strongly induced upon nitrogen depletion in RHA1 growing on benzoate, exhibiting the highest transcript levels of all 16 *atf* genes under nitrogen limiting conditions (12). In contrast, *atf6* transcripts were most abundant under nitrogen excess conditions with no apparent transcriptional induction upon nitrogen depletion. WS/DGAT genes identified in soil communities were exclusively from Actinomycetales (54), which includes the steroid-degrading suborder *Corynebacterineae*. In contrast, WS/DGAT genes from marine sediments were exclusively of gammaproteobacterial origin, predominantly from members of the oligotrophic marine Gammaproteobacteria (OMG) group, of which we recently identified several sterol-degrading strains (22, 23). BLASTp analysis (max E-value 10^-15^) of AtfA from *Acinetobacter calcoaceticus* ADP1 against the proteomes of the cholesteryl ester accumulating OMG bacteria *Zhongshania* sp. strain SB11-1A, *Haliaceae* sp. strain BC5-1 and the BD1-7 clade bacterium SB11-3 suggested that these strains encode between four and six WS/DGAT paralogs, further supporting our hypothesis that these proteins might function as steryl ester synthases.

Our results indicate that accumulated SEs can be reactivated and degraded in RHA1 upon relief of growth-limiting conditions, suggesting that SEs might play a vital role as a carbon and energy reservoir in these environments where fluctuating nutrient availability is common. We searched the genomes of strain RHA1 and of *Mtb* Erdmann for homologs of three prokaryotic and fifteen eukaryotic proteins with characterized or predicted sterol esterase activity available in the TrEMBL and Swiss-Prot databases (EC 3.1.1.13). However, none of the query proteins had significant similarity (accepted coverage > 85%, percent identity > 30%, E-values < 0.01) to RHA1 or *Mtb* proteins. Strain RHA1 encodes up to 54 putative lipases/esterase enzymes (33), which were suggested to contribute to neutral lipid degradation, but no distinct degradation system for endogenous TAGs or WEs has been identified so far. In contrast, the lipase family protein LipY in *M. tuberculosis* has been shown to have TAG degradation activity (55).

Interestingly, several animal lineages such as insects and soil-dwelling nematodes have lost the ability to synthesize sterols *de novo* (56) and thus rely on uptake of sterols with their diet (57). It is tempting to speculate that sterol and steryl ester accumulating bacteria in oligotrophic habitats such as soils or freshwater systems might constitute an important source for sterols for bacteriovorous, sterol-auxotrophic eukaryotes such as *Caenorhabditis elegans* or *Daphnia magna*.

RHA1 was also able to form esters of the phytosterols β-sitosterol, β-sitostanol, and campesterol under nitrogen-limiting conditions. In recent years, plant-derived phytosterols and their esters were recognized as potential agents to lower serum cholesterol levels in hypercholestemic patients (58, 59) by inhibiting absorption of endogenous, dietary cholesterol in the intestine. Today phytosterols and phytosteryl esters are approved in many countries as safe food additives, and several so-called functional foods are marketed worldwide, which are supplemented with phytosterol and their esters (60). Industrial production of phytosteryl esters is mainly accomplished by chemical esterification or transesterification of plant-derived sterols, which requires high temperatures and the use of strong acids or bases (61). Our study provides a basis for biotechnological production of phytosteryl esters using non-pathogenic whole-cell bacterial systems, thus eliminating the requirement for elevated temperature and acid or base catalysts. Similar to RHA1, many other sterol-degrading Actinobacteria are also able to use phytosterols as growth substrates, providing an excellent screening resource for efficient phytosteryl ester accumulation.

## MATERIAL AND METHODS

### Bacteria and culture conditions

*Rhodococcus jostii* strain RHA1, as well as *Mycobacterium smegmatis* strain mc^2^155, *Rhodococcus opacus* strain PD630, *Rhodococcus ruber* strain Chol-4, and *Amycolatopsis* sp. strain ATCC 39116 were cultivated in M9 minimal medium supplemented with trace elements and vitamin B1 at 30 °C. The marine bacteria *Mycobacterium* strain BC8-1, *Zhongshania* sp. strain SB11-1A, *Haliaceae* sp. strain BC5-1, and the BD1-7 clade bacterium strain SB11-3 were cultivated in artificial marine medium at 21 °C as described earlier (23). Substrates were added at the following concentrations: cholesterol 1 mM, sodium palmitate 1.0 mM or 1.7 mM, succinate 7.0 mM, glucose 5 mM. Starter cultures were grown on 10 mM succinate or 1 mM cholesterol. Cholesterol and other sterol substrates were added as solids and 1% (w/v) methyl-β-cyclodextrin was added to the medium, when required, prior to autoclaving. Substrate removal was monitored by gas chromatography-coupled mass spectrometry (GC-MS) of organic extracts of acidified culture supernatants as described earlier (62). Nitrogen-limiting conditions were achieved by reducing the concentration of ammonium chloride in the medium from 18.70 mM to 0.93 mM. Under N-excess conditions the ratio of carbon to nitrogen in the growth medium was around 1.5:1, while it was 29:1 under N-limiting conditions. To limit nitrogen transfer between cultures, starter cultures were washed twice with phosphate-buffered saline (PBS) without nitrogen. Growth was measured as protein, using the bicinchoninic acid assay (Thermo Scientific) after hot alkaline cell lysis.

*Mycobacterium tuberculosis* strain Erdmann (*Mtb*) was grown in 7H9 medium supplemented with 0.5% tyloxapol and 0.5 mM cholesterol at 37 °C. If required 0.5 mM sodium palmitate were spiked into growing cultures after 5 days. Growth was measured as optical density at 600 nm. To test effects of nitric oxide (NO) stress and iron limitation, cholesterol-grown *Mtb* was harvested and washed in 7H9 medium without carbon source, and cells were suspended in the same medium containing 0.5 mM cholesterol and/or 0.5 mM palmitate. 0.5 mM of the NO releasing agent spermine-NONOate was added from a 10 mM stock in 0.01 M NaOH at the beginning of the experiment and again after 16 h of incubation. Spermine was added at the same concentration to control cultures. The iron chelating agent deferoxamine was added at 0.5 mM from a 10 mM stock at the beginning of the experiment.

Biosample IDs and references to cholesterol degradation capability of all strains used in this study are summarized in **supplemental Tab. S1**.

### Ammonium quantification

Ammonium concentrations were quantified using indophenol blue. Culture supernatant samples were diluted 25- to 500-fold in distilled water and 200 µl aliquots were distributed into 96-well plates in triplicates. 8 µl of reagent (10% w/v phenol in ethanol), 8 µl catalyst (0.5% w/v nitroprusside in water) and 20 µl oxidizing solution (20% w/v trisodium citrate and 1% w/v sodium hydroxide solution, mixed with 5% sodium hypochlorite 4:1 v/v) were added and plates were incubated at 37 °C for 1-2 h in the dark. Absorbance was measured at 630 nm. Ammonium concentrations were calculated using a standard curve with 0-2 mg ml^-1^ ammonium chloride. Alternatively, semi-quantitative ammonium test strips (Quantofix®, Macherey-Nagel, Germany) were used for instant ammonium quantification.

### Lipid extraction and analysis

For lipid extraction, cells were harvested, washed twice with distilled water and frozen at - 80 °C. *Mtb* cells were autoclaved before freezing. Cell pellets were lyophilized and their dry weight was determined by weighing the pellets. The pellets were then suspended in 1 ml distilled water and broken by bead beating (four rounds of 60 sec at 4.5, 5.0, 5.5, and 6.0 speed setting, respectively, in a FastPrep-24 bead beater; MP Biomedicals, Solon, OH). Between rounds samples were incubated on ice for 5 min. Neutral lipids were extracted with chloroform-methanol (2:1, v/v, 1% acetic acid), dried under nitrogen and suspended in chloroform with 1% acetic acid. If required cholestane was added as internal standard from a 10 mM stock solution in chloroform prior to organic extraction for normalization and quantification of cholesterol and FAMEs. Aliquots of neutral lipid extracts were analyzed by thin layer chromatography (TLC) on 60 F_254_ silica gel plates (Merck) using hexane/diethyl-ether/acetic-acid (90:8:1, v/v/v) as a solvent system. Cholesterol palmitate, cholesterol, and tripalmitin dissolved in chloroform were used as standards. The plates were dried, dabbed with 10% cupric sulfate in 8% phosphoric acid solution and developed at 200 °C for 5-10 min. Native neutral lipid extracts from non-lyophilized cell pellets of strain RHA1 grown under nitrogen limiting conditions with cholesterol or succinate were also analyzed using a GC-coupled flame ionization detector (FID). For this, 20 ml culture aliquots were harvested, washed twice and frozen at −80 °C at different time points after nitrogen was depleted from the medium. Neutral lipids were extracted with tert-butyl ether:methanol (10:3, v/v), followed by two extractions with hexane. Combined organic extracts were dried under nitrogen and suspended in chloroform. Neutral lipid extracts were analyzed by gas chromatography on an Agilent 8890 series GC system equipped with a J&W Select Biodiesel GC Column (15 m, 0.32 mm, 0.10 µm), coupled to a flame ionization detector (FID). GC-injector and detector temperatures were set to 350 °C, and 400 °C, respectively. 1 µl sample volumes were injected with an initial purge flow rate of 50 ml min^-1^, which subsequently decreased to 20 ml min^-1^ after 2 min, and a septum purge flow rate of 3 ml min^-1^. The helium flow rate was 8.09 ml min^-1^. The oven was set to an initial temperature of 40 °C for 2 min with an equilibration time of 0.5 min. The temperature increased by 25 °C min^-1^ and was held at 170 °C for 2 min, then increased by 15 °C min^-1^ to 210 °C, then by 5 °C min^-1^ to 250 °C and held for 5 min. The temperature finally increased by 10 °C min^-1^ to 400 °C and held for 2 min. The FID gas flow settings were set to 400 ml min^-1^ air flow, 30 ml min^-1^ H_2_ fuel flow and 25 ml min^-1^ makeup flow (N_2_). Cholesteryl palmitate, cholesteryl heptadecanoate, and tripalmitin TAG dissolved in chloroform were used as standards. CE and TAG peaks were normalized with the internal standard cholestane for quantification.

### Isolation of lipid droplets

Lipid droplets were isolated as previously described (63, 64). Briefly, RHA1 cultures were harvested by centrifugation (4,000 x g for 10 min), washed three times with 40 ml Buffer A (25 mM tricine, 250 mM sucrose, pH 7.8) and suspended in 8 ml buffer A. After incubation on ice for 20 min, cells were disrupted by passing them three times through a cooled French pressure cell at 100 MPa. Cell debris was removed by centrifugation at 4,000 × g for 10 min. Supernatants were loaded into SW40 tubes (Beckman), layered with 2 ml Buffer B (20 mM HEPES, 100 mM KCl, 2 mM MgCl2, pH 7.4), and centrifuged at 180,000 x g for 1 h at 4 °C. The lipid droplet fraction on top of the sucrose gradient was transferred to a 1.5 ml Eppendorf tube and was washed twice with 200 μl Buffer B and finally suspended in 1 ml buffer B. No pellets were formed during washing of the lipid droplets suggesting that lipid droplets were not contaminated by membrane particles. Total lipids were extracted with 5 ml chloroform-acetone (1:1, v/v). The organic phase was dried under nitrogen and lipids were suspended in 1 ml chloroform with 1% acetic acid. Aliquots were analyzed by TLC as described above.

### Lipid fractionation

For analyzing the fatty acid profile of cellular lipids and lipid droplets, lipid extracts were fractionated stepwise. Total lipid extracts derived from lipid droplets were first run over a silica column equilibrated with chloroform and neutral lipid fractions were obtained by eluting with 6 column volumes chloroform with 1% acetic acid. Glycolipid fractions were subsequently eluted with 6 column volumes acetone (not analyzed further) followed by phospholipid elution with 7.5 column volumes methanol. Neutral lipids from lipid droplets and cellular lipid extractions were further fractionated by silica column chromatography. Neutral lipid samples were dried, resuspended in hexane and poured on a silica column equilibrated with the same solvent. 5 different fractions were obtained by eluting sequentially with 5 column volumes hexane, 6 column volumes hexane/diethyl ether (99:1, v/v), 5 column volumes hexane/diethyl ether (95:5, v/v), 5 column volumes hexane/diethyl ether (92:8, v/v), and 6 column volumes chloroform with 1% acetic acid. Fractions were dried under nitrogen and lipids were suspended in 1 ml chloroform with 1% acetic acid. If required, aliquots were analyzed by TLC as described above.

### Alkaline transesterification with methanol and lipid quantification

Dried cellular lipid extracts or individual lipid fractions were subjected to alkaline transesterification with methanol modified after (65). For this, dried lipids were resuspended in methanol/toluene (1:1, v/v), mixed with an equal volume of methanolic KOH and incubated for 1 h at 50 °C. Phospholipid fractions were incubated for 15 min at 30 °C. Free sterols and fatty acid methyl-esters were extracted twice with 4 volumes hexane/chloroform (4:1, v/v) from 0.15 volumes acetic acid (1 M) and 1 volume water. The organic phase was dried under nitrogen and lipids were suspended in 1 ml chloroform with 1% acetic acid. Aliquots were analyzed by GC-MS as described earlier for FAMEs and free cholesterol (62) and for wax esters (32). Cholesterol and fatty acid methyl-esters (FAMEs) were identified by comparison to authentic standards as well as matching against the NIST database (National Institute of Standards and Technology, v11). To quantify total neutral lipids, unfractionated neutral lipid extracts were subjected to transesterification and FAMEs were normalized on cholestane. To quantify SEs in cellular lipid extracts, fractions 1 (containing the internal standard cholestane) and 2 (containing steryl esters) were combined prior to transesterification. This allowed normalisation and quantification of free cholesterol derived from transesterification. The percentage of CEs per extracted cellular dry weight was estimated by calculating the total weight of CEs from CE containing fractions and dividing it by the dry weight of the respective extracted cell pellet. For this, the cholesterol concentration in transesterified fractions was calculated using a cholesterol-cholestane standard curve. Subsequently, this was transformed into the total weight of CEs using the molecular masses of all detectable CEs weighted on the peak area ratio of the respective detected FAMEs (C15:0, C16:1, C16:0, C17:1, C17:0, C18:1, C18:0). To estimate ratios of neutral lipid classes in lipid droplets, cholestane and methyl-nonadecanoate were added as internal standards to each fraction prior to transesterification from 10 mM stock solutions in chloroform. Free cholesterol (in fraction 5) and cholesterol released from transesterification of fraction 2 were normalized on cholestane and FAMEs derived from transesterification of CEs and TAGs were normalized on methyl-nonadecanoate before calculating their molar ratios.

### Fluorescence microscopy

To localize accumulating neutral lipids in strain RHA1, we added the fluorescent dye Nile Red to growing cholesterol cultures under N-limiting or N-excess conditions without cyclodextrin. Nile red was added to the medium after autoclaving at a final concentration of 0.5 µg ml^-1^ from a sterile stock solution in DMSO (0.3 mg ml^-1^). Nitrogen and cholesterol depletion were confirmed as described above. Cells from both cultures were examined four days after reaching stationary phase. Cells were washed twice and resuspended in sterile PBS before microscopy. Slides were examined on a Zeiss Axio Imager M1 fluorescence microscope equipped with a Zeiss HBO 103 W/2 high-pressure mercury lamp and an AxioCam MRm camera in phase contrast, differential interference contrast or fluorescence mode. Fluorescence images were shot using excitation and emission wavelengths of 550 nm and 605 nm, respectively. All Images were processed using the Zeiss AxioVision software (40 V 4.6.3.0).

To further analyze the localization of accumulating cholesterol within cells of RHA1 under nitrogen-limiting conditions, we performed a cell suspension experiment with fluorescently labeled cholesterol (cholesterol-BODIPY) as substrate under N-limiting conditions followed by laser scanning confocal microscopy. For that, cholesterol-grown cells were washed two times with PBS without nitrogen. Depletion of cholesterol in the starter culture was confirmed by organic extraction and GC-MS analysis. Cells were suspended in 1/5 of the original volume of M9 medium without nitrogen containing 0.5 mM palmitate, 0.5 mM fluorescently labeled cholesterol (cholesterol-BODIPY) and 1% (w/v) methyl-β- cyclodextrin and incubated for 48 h at 200 rpm and 30 °C. Cells were washed twice with PBS without nitrogen containing 0.02% (w/v) cholate and 0.02% (v/v) tylaxopol and suspended in PBS. Cells were analyzed using a laser scanning confocal microscope (Olympus Fluoview FV1000-Inverted). A small aliquot of cells was broken by ultrasonication (four rounds of 20 sec using a Microsom cell disruptor XL set to 5; Mandel scientific Co. Ltd., Guelph, ON, Canada) and neutral lipid extracts were analyzed by TLC. Undeveloped plates were analyzed using a GE Typhoon trio + imager with excitation wavelength of 488 nm and an emission filter of 520 nm. Pre-tests with unlabelled cholesterol and palmitate showed that CE accumulation in RHA1 cell suspensions started within 22 h of incubation and continued at least until 95 h of incubation (data not shown).

## Acknowledgments

We thank Fanrui Meng and Carl Strittmatter for help with fluorescence microscopy. This study was funded by a Discovery Grant from the Natural Sciences and Engineering Research Council of Canada.

## Data availability

Draft genomes of *Mycobacterium* strain BC8-1, *Zhongshania* sp. strain SB11-1A, *Haliaceae* sp. strain BC5-1, and the BD1-7 clade bacterium strain SB11-3 have been deposited in the European Nucleotide Archive (ENA) under the BioProject number PRJEB30766.

