## Supplemental material for "Steryl ester formation and accumulation in steroid-degrading bacteria"

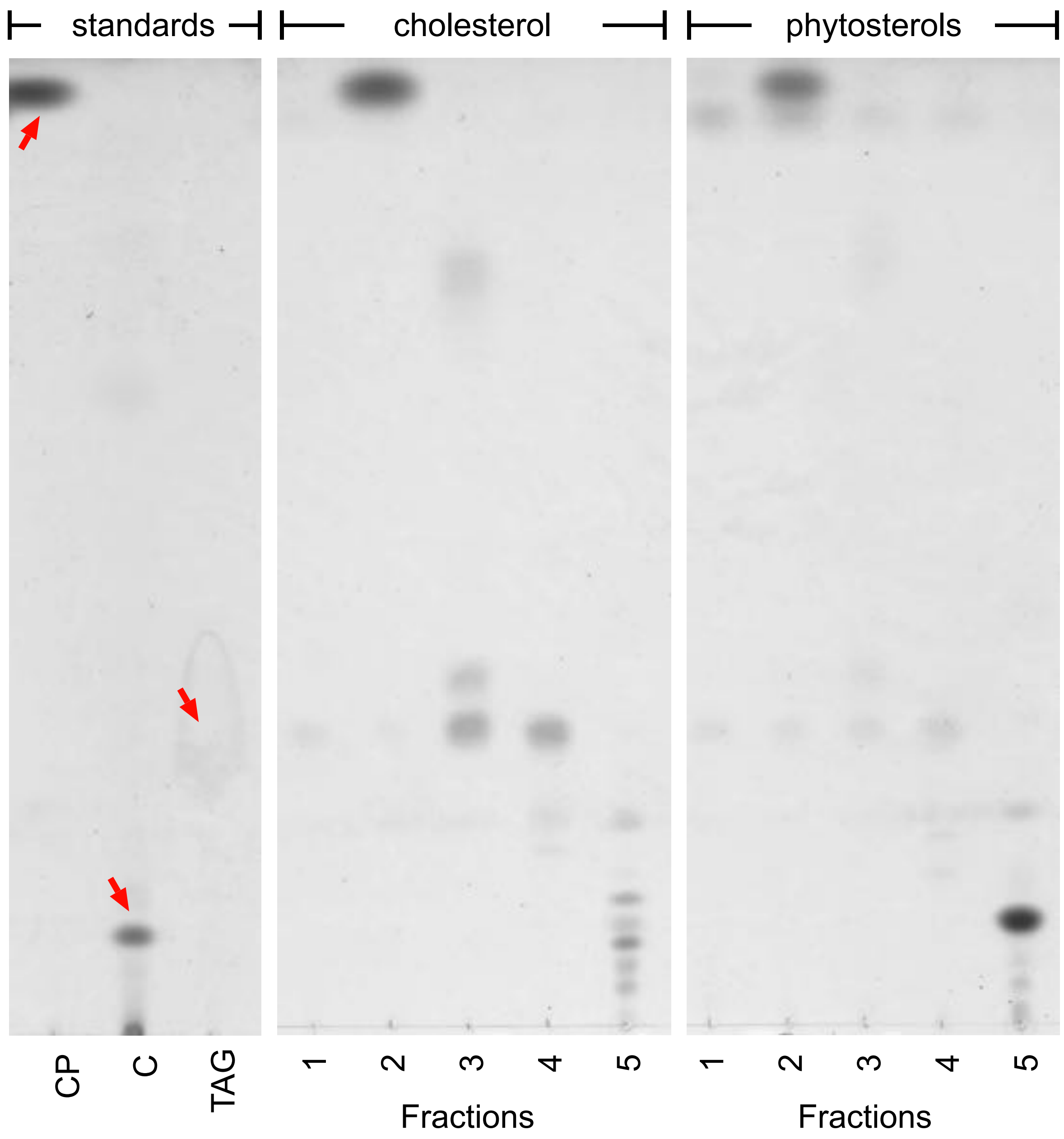


**Supplemental Fig. S1:** TLC analysis of fractionated neutral lipid extracts of RHA1 cells grown with cholesterol or phytosterols. SEs eluted in fraction 2. Red arrows indicate standard spots. Abbreviations: CP, cholesterol palmitate; C, cholesterol, TAG, triacylglycerol.

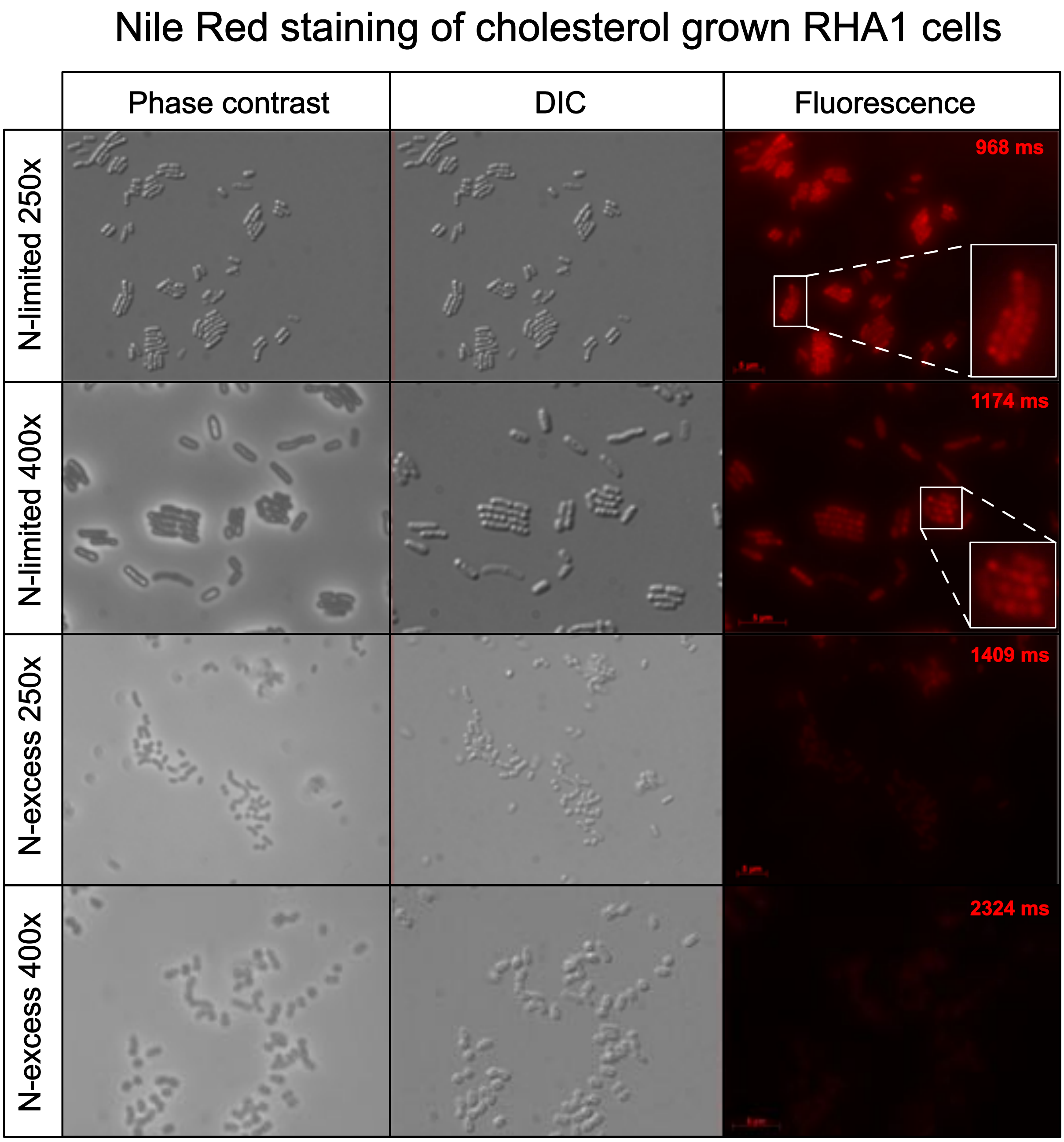


**Supplemental Fig. S2:** Microscopic analysis of *Rhodococcus jostii* RHA1 grown with cholesterol under N-limiting or N-excess conditions in the presence of the fluorescent dye Nile Red. Cells from N-limited and N-excess cultures were examined four days after reaching the stationary phase. Slides were examined on a Zeiss Axio Imager M1 fluorescence microscope and an AxioCam MRm camera in phase contrast, differential interference contrast (DIC) or fluorescence mode. Fluorescence images were shot using excitation and emission wavelengths of 550 nm and 605 nm respectively. All Images were processed using the Zeiss AxioVision software (40 V 4.6.3.0). Scale bars = 5 µm. Large white boxes show magnification of the small boxed areas in fluorescence micrographs for better visualization of the accumulating lipid droplets.

Cholesterol-BODIPY staining of RHA1 cells

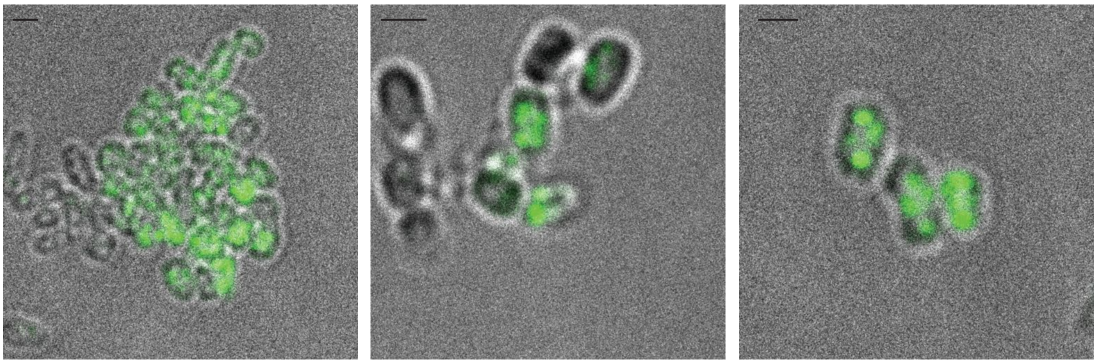

**Supplemental Fig. S3:** Confocal microscopy images of *Rhodococcus jostii* RHA1 cells incubated with fluorescently labelled cholesterol (cholesterol-BODIPY) plus palmitate for 48 h without a nitrogen source. Scale bar = 1 µm. Excitation/emission wavelengths 495 nm/508 nm.

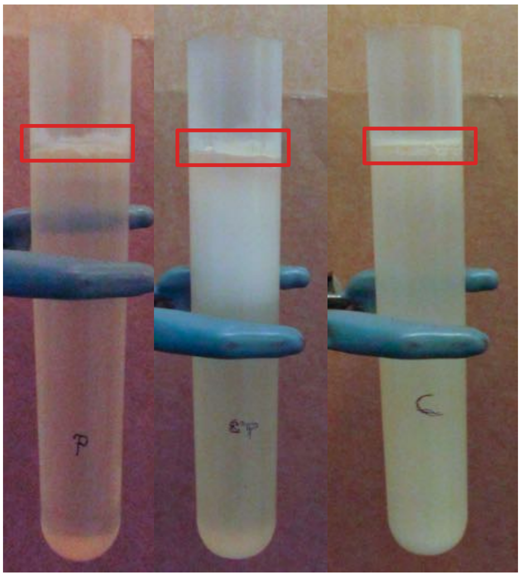


**Supplemental Fig. S4:** Lipid droplets on top of sucrose gradient isolated from *Rhodococcus jostii* RHA1 grown on palmitate, palmitate plus cholesterol or cholesterol under N-limiting conditions.

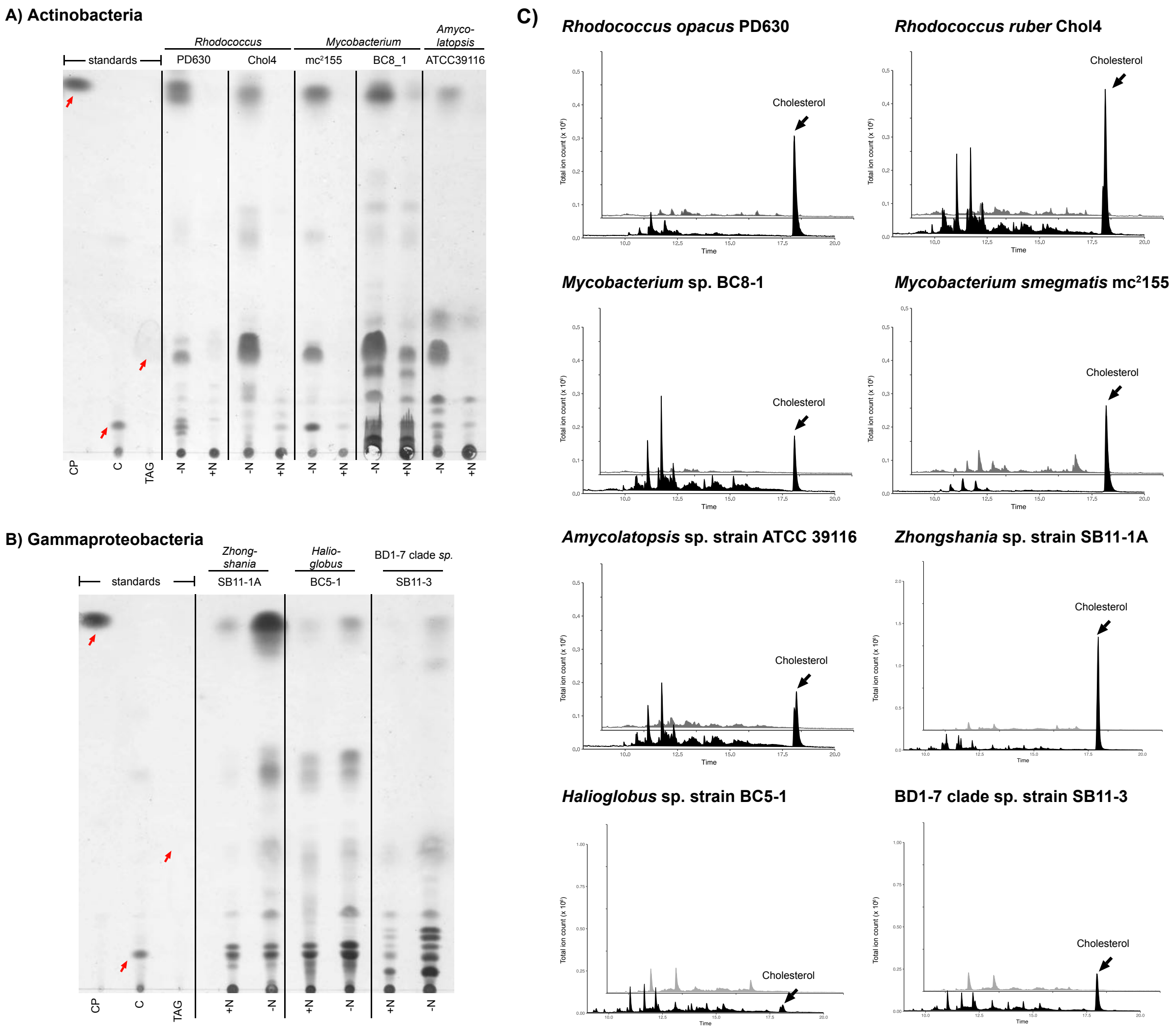


**Supplemental Fig. S5 (A)** TLC analysis of neutral lipid extracts from *Rhodococcus opacus* PD630, *Rhodococcus ruber* Chol4, *Mycobacterium* sp. BC8-1, *Mycobacterium smegmatis* mc$^2$155, and *Amycolatopsis* sp. ATCC 39116 cells grown on cholesterol under N-limiting (-N) and N- excess conditions (+N) in stationary phase. **(B)** TLC analysis of neutral lipid extracts from *Zhongshania* sp. strain SB11-1A, *Halioglobus* sp. strain BC5-1, and BD1-7 clade sp. strain SB11-3 cells grown on cholesterol under N-limiting (-N) and N-excess conditions (+N) in stationary phase. Red arrows indicate standard spots. Abbreviations: CP, cholesterol palmitate; C, cholesterol; TAG, triacylglycerol. **(C)** GC-MS analysis of transesterified Fraction 2 (black) and untreated Fraction 2 (grey) of neutral lipid extracts from N-limited cultures.

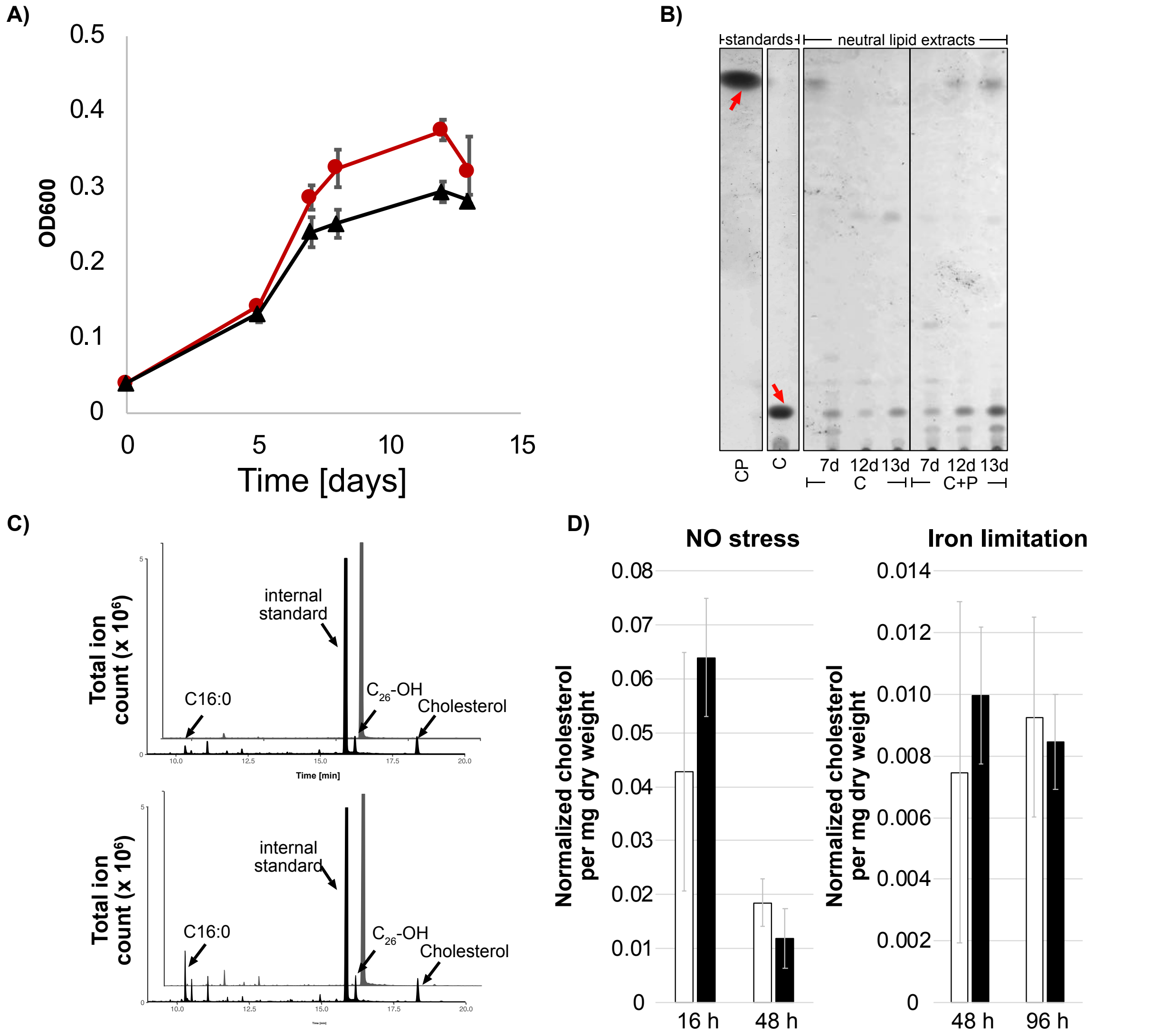


**Supplemental Fig. S6 (A)** Growth of *Mycobacterium tuberculosis* (*Mtb*) Erdmann on 0.5 mM cholesterol (red) or 0.5 mM cholesterol plus 0.5 mM palmitate added on day 5 (black); n = 3. **(B)** TLC analysis of neutral lipid extracts of *Mtb* cells grown on cholesterol (C) or cholesterol plus palmitate (C+P). Red arrows indicate standard spots. Abbreviations: CP, cholesterol palmitate; C, cholesterol. The volume of neutral lipid extracts applied to TLC plates was normalized on cellular dry weight of the extracted cultures. TLC plates were analyzed for three replicate cultures and one of these plates is shown here. Replicate cultures showed same results. **(C)** GC-MS analysis of transesterified Fraction 2 (black) and untreated Fraction 2 (grey) of neutral lipid extracts from a cholesterol-grown culture harvested on day 7 (top) and of a cholesterol plus palmitate-grown culture harvested on day 13 (bottom). **(D)** Quantification of accumulating CEs in *Mtb* cell suspensions containing cholesterol plus palmitate treated with nitric oxide (NO) or the iron chelator deferoxamine (black bars) and control cultures without NO or deferoxamine (white bars) expressed as normalized cholesterol per mg dry weight (n = 3).

**Supplemental Tab. S1:** Biosample IDs and references to sterol degradation ability of all strains used in this study

| | Biosample ID | Reference to sterol degradation |
|---|---|---|
| *Rhodococcus jostii* strain RHA1 | SAMN02604146 | Van der Geize R, et al. 2007. PNAS 104(6): 1947-1952. https://doi.org/10.1073/pnas.0605728104 |
| *Mycobacterium smegmatis* strain mc$^2$155 | SAMN02603392 | Uhía I , et al. 2012. Env Microbio Reports, 4: 168-182. https://doi.org/10.1111/j.1758-2229.2011.00314.x |
| *Rhodococcus opacus* strain PD630 | SAMN02821848 | Holder JW, et al. 2011. PLoS Genet 7(9): e1002219. https://doi.org/10.1371/journal.pgen.1002219 |
| *Rhodococcus ruber* strain Chol-4 | SAMN01804522 | Fernández de las Heras L, et al. 2009. Curr Microbiol 59: 548. https://doi.org/10.1007/s00284-009-9474-z |
| *Amycolatopsis* sp. strain ATCC 39116 | SAMN00117521 | Bergstrand LH, et al. 2016. mBio 7(2):e00166-16. https://doi.org/10.1128/mBio.00166-16. |
| *Mycobacterium* strain BC8-1, | SAMEA5215812 | Holert J, et al. 2018. mBio 9:e02345-17. https://doi.org/10.1128/ mBio.02345-17. |
| *Zhongshania* sp. strain SB11-1A | SAMEA5215815 | Holert J, et al. 2018. mBio 9:e02345-17. https://doi.org/10.1128/ mBio.02345-17. |
| *Haliaceae* sp. strain BC5-1 | SAMEA5215810 | Holert J, et al. 2018. mBio 9:e02345-17. https://doi.org/10.1128/ mBio.02345-17. |
| BD1-7 clade bacterium strain SB11-3 | SAMEA5215816 | Holert J, et al. 2018. mBio 9:e02345-17. https://doi.org/10.1128/ mBio.02345-17. |
| *Mycobacterium tuberculosis* strain Erdman | SAMD00061050 | Crowe AM, et al. 2017. mBio 8:e00321-17. https:// doi.org/10.1128/mBio.00321-17. |
